# “Metabolic dysregulations of cancer cells with metastatic potential”

**DOI:** 10.1101/2021.06.02.446725

**Authors:** Sara Abdul Kader, Shaima Dib, Iman W. Achkar, Gaurav Thareja, Karsten Suhre, Arash Rafii, Anna Halama

## Abstract

Metastasis is the primary cause of cancer related deaths due to the limited number of efficient druggable targets. Signatures of dysregulated cancer metabolism could serve as a roadmap for the determination of new treatment strategies, given their vital role in cancer cell responses to multiple challenges, including nutrient and oxygen availability. However, the metabolic signatures of metastatic cells remain vastly elusive. We conducted untargeted metabolic profiling of cells and growth media of five selected triple negative breast cancer cell lines with high metastatic potential (HMP) (MDA-MB-231, MDA-MB-436, MDA-MB-468) and low metastatic potential (LMP) (BT549, HCC1143). We identified 92 metabolites in cells and 22 in growth medium that display significant differences between LMP and HMP. The HMP cell lines had elevated level of molecules involved in glycolysis, TCA cycle and lipid metabolism. We identified metabolic advantages of cell lines with HMP beyond enhanced glycolysis by pinpointing the role of branched chain amino acids (BCAA) catabolism as well as molecules supporting coagulation and platelet activation as important contributors to the metastatic cascade. The landscape of metabolic dysregulations, characterized in our study, could serve in the future as a roadmap for the identification of treatment strategies targeting cancer cells with enhanced metastatic potential.

## 1. Introduction

Metastatic disease accounts for approximately 90% of cancer related deaths [1,2], despite relatively low metastatic efficiency due to the challenging multistep cascade required to establish colonies in distant tissue [3]. Triple negative breast cancer (TNBC), characterized by the lack of expression of the estrogen receptor (ER), progesterone receptor (PR) and human epidermal growth factor receptor 2 (HER2) [4], tends to display a more aggressive clinical course with frequent distant recurrence and thus poor prognosis compared to other breast cancer types [5]. Lack of available targeted therapy for TNBC patients along with the limited understanding of the molecular processes governing metastatic disease reflect on very narrow treatment options for those patients [6]. Therefore, further insights into molecular events related to metastasis could revel novel treatment targets.

Metabolomics, provides almost unbiased overview of the current processes that are ongoing in the biological system by monitoring the levels of endogenous and exogenous small molecules (metabolites) in that system [7]. Hence, metabolic profiling can precisely inform on altered molecular pathways and responses to environmental stimuli. The metabolic signatures discriminating healthy from disease are frequently deployed for biomarkers identification but also to provide insights into the pathological processes causing disease [8–10]

In the last decade, our view on cancer as being a strictly genetic disease has evolved and nowadays, cancer is also considered as a metabolic disorder [11]. This insight arises from the vast body of evidence from multiple studies showing drastic differences between metabolism of cancer and normal cells in glycolysis, glutaminolysis, nucleotide metabolism, as well as synthesis and catabolism of lipids [12–16]. The metabolic dysregulations observed in cancer cells can became a basis for new drug discoveries [10], designed to target cancers with e.g. enhanced glutaminolysis [17] and fatty acid synthesis [18], as well as for the identification of cancer survival mechanisms under treatment [19–21]

Recently, cancer metabolic plasticity related to the ability of cancer cells to fulfill the metabolic requirements of the metastatic cascade, as well as metabolic flexibility related to cancer cells’ use of different nutrients to meet the energetic requirements during metastasis, were defined as key contributors enabling cancer cell adjustment during metastasis [22]. The impact of metabolic rewiring on metastatic signaling cascade was also suggested [23]. For instance, dysregulations in TCA cycle metabolism towards accumulation of fumarate [24] and succinate [25], as well as the generation and accumulation of 2-hydroxyglutarate [26,27] were linked to DNA methylation and associated with epithelial-mesenchymal transition (EMT) [28]. The role of lipid metabolism as well as glycosylation were defined by us and others as important steps in nesting of cancer cells in the endothelial niche [29,30]. Alterations in metabolism of acetyl-CoA, recognized as an epigenetic regulator for its involvement in histone acetylation, were identified as important components of EMT [31]. However, the metabolic program related to metastatic potential of cancer cell remains largely elusive.

Here, we investigated whether TNBC cell lines harboring different metastatic potential *in vivo* would differ metabolically *in vitro.* To this end, we selected five TNBC cell lines (BT549, HCC1143, MDA-MB-231, MDA-MB-436, and MDA-MB-468), for which metastatic potential was recently defined by Jin et al. [32], who created a metastasis map (MetMap), by characterizing organ-specific patterns of metastasis and metastatic potential of 500 different human cancer cell lines from 12 different types of solid tumors, including breast cancer [32]. We used untargeted metabolomics profiling to describe metabolism of TNBC cell lines defined as with low (BT549 and HCC1143) and high (MDA-MB-231, MDA-MB-436, and MDA-MB-468) metastatic potential along with normal epithelial breast cell line (hTERT-HME1). We detected 479 metabolites, which allowed a clear separation between normal and TNBC cell lines on the principal component analysis (PCA) score plot. A total of 291 metabolites displayed significant differences at a false discovery rate (FDR) < 0.01 between normal and TNBC cell lines. Next, we searched for metabolic differences discriminating cell lines with low metastatic potential (LMP) from those with high metastatic potential (HMP), in both, cultured media and cells extracts. We found that cell lines with LMP and HMP are metabolically different, and those differences are independent of canonical EMT markers. We identified enrichment in glycolysis and citrate metabolism as well as enhanced BCAA catabolism and dysregulated metabolism of lipids as signatures of HMP cell lines. Additionally, we found metabolic features potentially involved in coagulation and platelet activation in HMP cell lines. Our findings shed new light on the landscape of metabolic dysregulations related to the metastatic potential of TNBC cell lines, which could in the further be considered as therapeutic targets.

## 2. Materials and Methods

### 2.1. Culture conditions

The established cancer cell lines (BT-549, HCC1143, MDA-MB-231, MDA-MB-436, MDA-MB-468) all TNBC models and the normal epithelial breast cancer cell line (hTERT-HME1 [ME16C]) were purchased from American Type Culture Collection (ATCC, Manassas, VA, USA). All TNBC cell lines were maintained at 37°C and 5 % CO2 and grown in Roswell Park Memorial Institute medium (RPMI-1640) media supplemented with 10% fetal bovine serum and 1% penicillin-streptomycin. The growth media of BT-549 was supplemented with 1 μg/ml insulin and growth media of MDA-MB-436 with 10 μg/ml insulin and 16 μg/ml glutathione, as per ATCC instructions.

The cell lines dedicated for metabolomics and western blot analysis were cultured and prepared at the same time points on separate Petri dishes with a growth area of 21 cm^2^. The study was conducted in two independent experiments each conducted in triplicates. The cell lines MDA-MB-231, MDA-MB-436, MDA-MB-468 were seeded at the density of 1×10^6^ and BT-549 and HCC1143 at the density 1.5×10^6^ per dish. 24h after seeding, the medium was changed with fresh medium and cells were incubated for an additional 24h. At the day of collection, the cells reached around 85 % of confluency. The collection process for metabolomics and for Western blot was conducted 48h after seeding and the description is provided in section 2.2 and 2.3, respectively.

### 2.2. Western blot

The medium was aspirated, the cells were washed with phosphate buffered saline (PBS) and incubated for around 1 min with 1 mL of trypsin at 37°C in the incubator. 1.5 mL media was added to the cells resuspended and placed into a 15 mL tube. The samples were centrifuged for 5 min at 400 x g, the supernatant was removed, and the cells were resuspended in PBS. The samples were centrifuged, the supernatant was removed, and the cell pellets placed at −80°C until further processing.

At the day of processing, the samples were thawed on ice and mixed at the ratio of 1×10^6^ cells/50 μL with lysis buffer Nonidet P-40 (NP40) supplemented with protease-phosphatase cocktail inhibitors and phenyl methane sulfonyl fluoride (PMSF). The samples were lysed by three freeze-thaw cycles as previously described [33]. The supernatant was collected after centrifugation for 10 min at 18,000 x g. The total protein content was quantified using the DC protein assay kit (Bio-Rad, Richmond, CA). The proteins were denatured by incubation with 1 × Laemmli buffer containing β-mercaptoethanol, at 95 °C for 10 min.

The prepared whole cell lysates were used to conduct gel electrophoresis followed by transfer to a polyvinylidene fluoride membrane (Bio-Rad). The membrane was blocked in 5% milk solution in Tween-PBS (PBS with 0.1% Tween 20) for 1 h followed by overnight incubation in a primary antibody at 4 °C. The membrane was washed three times in Tween-PBS and incubated in the corresponding secondary antibody for 1h at room temperature. The membrane was washed three times prior to development. Both primary and secondary antibodies were prepared in the recommended dilution in a 5% milk or bovine albumin solution in Tween-PBS. The signal was detected by a chemiluminescent western blot detection kit (Thermofischer) and the blots were developed and visualized under a ChemiDoc system (Amersham, Bio-Rad, USA). The primary antibodies used were Twist2 (GeneTex, #GTX50850), MMP-2 (Cell signaling, #40994), Vimentin (Cell signaling, #5741), N-Cadherin (Cell signaling, #13116), E-Cadherin (Cell signaling, #14472), beta-Tubulin (Cell signaling, #2146s), and beta-Actin (Cell signaling, #3700S). The corresponding secondary antibodies included horseradish peroxidase–conjugated anti-mouse (Cell Signaling) and anti-rabbit (Cell Signaling).

### 2.3. Sample preparation for metabolic analysis

The growth media was collected into the collection tube, centrifuged for 5 min at 400 g, 500 μL was placed into fresh collection tube and flash frozen in liquid nitrogen. The samples were stored at −80°C until shipment.

The cell processing for metabolic analysis was conducted as previously described [34]. Briefly, the cells were washed twice with 37 °C PBS. 1 mL of ice-cold 80% methanol in H_2_O was added per dish, and the cells were scraped off from the dish. The scraped-in-methanol cells were placed in a collection tube and flash-frozen in liquid nitrogen, and stored at −80 °C until further processing. Metabolite extraction out of the cells was conducted in a series of three freeze-thaw cycles; the samples were thawed on ice for 5 min followed by freeze in liquid nitrogen for 5 min. The samples were centrifuged at 18,000 x g for 5 min at 4°C, transferred to a fresh collection tube and stored at −80°C until shipment.

The remaining pellets were used for the determination of protein content in the samples to account for differences in cell growth. Sample processing was conducted as previously described [35]. Briefly, the remaining pellets were dried in a speed vacuum for 20 min. 60 μL of 0.2 M NaOH was added into the dried samples heated for 20 min at 95 °C with frequent vortexing. The samples were centrifuged at 18,000 × g for 5 min and the supernatant was transffered to a fresh collection tube. Protein content was determined using the Bio-Rad DC protein assay, relative to bovine serum albumin standards (0–1.8 mg/mL).

The growth media and cell extract were shipped to Metabolon Inc. (Durham, NC, USA) on dry ice for metabolite measurements.

### 2.4. Metabolic measurements

Metabolic profiling of growth media and cell extracts was performed using Metabolon platforms deploying Waters ACQUITY ultra-performance liquid chromatography (UPLC) and a Thermo Scientific Q-Exactive high-resolution/accurate mass spectrometer interfaced with a heated electrospray ionization (HESI-II) source and Orbitrap mass analyzer, as previously described [36]

Proteins were precipitated from 100 μL of growth media with methanol using an automated liquid handler (Hamilton LabStar). The precipitated extract from growth media and cell extracts were split into four aliquots to undergo the following processes: 1) two fractions for analysis by two separate reverse-phase (RP)/UPLC-mass spectrometry (MS)/MS methods with positive ion mode electrospray ionization (ESI); 2) one fraction for analysis by RP/UPLC-MS/MS with negative ion mode ESI; 3) one fraction for analysis by hydrophilic interaction chromatography (HILIC)/UPLC-MS/MS with negative ion mode ESI. The samples were dried under nitrogen flow.

Dried samples were reconstituted in solvents compatible with each of the four methods: 1) acidic positive ion (optimized for hydrophilic compounds)—extract gradient eluted from a C18 column (Waters UPLC BEH C18–2.1 × 100 mm, 1.7 μm) with water and methanol containing 0.05% perfluoropentanoic acid and 0.1% formic acid; 2) acidic positive ion (optimized for hydrophobic compounds)—extract gradient eluted from C18 (Waters UPLC BEH C18–2.1 × 100 mm, 1.7 μm) with methanol, acetonitrile, water, 0.05% perfluoropentanoic acid, and 0.01% formic acid; 3) basic negative ion—extract gradient eluted from a separate dedicated C18 column using methanol and water containing 6.5 mM ammonium bicarbonate at pH 8; and 4) negative ionization—extract gradient eluted from a HILIC column (Waters UPLC BEH Amide 2.1 × 150 mm, 1.7 μm) using water and acetonitrile with 10 mM ammonium formate at pH 10.8. In the MS analysis, the scan range varied between methods but covered the range of 70–1000 m/z.

The raw data were extracted using Metabolon’s hardware and software. Compound’s identification was conducted by comparison of peaks to library entries of purified standards based on retention index, with an accurate mass match to the library of ±10 ppm, and MS/MS forward and reverse scores between the experimental data and authentic standards. The data was manually curated. The resulted metabolic data was normalized to correct variations resulting from inter-day tuning differences in the instrument. Each compound was corrected in a run-day. The metabolomics data is provided in Supplementary Table 1.

### 2.5. Statistical data analysis

The statistical data analysis was conducted using MetaboAnalyst 5.0 (https://www.metaboanalyst.ca/home.xhtml), a web server designed for comprehensive metabolomic data analysis, visualization and interpretation [37]. The normalized per run day metabolite intensities were further normalized by the sample protein content and submitted for analysis. The missing values were imputed by the min and the data was log scaled. The log scaled metabolite intensities were analyzed using parametric test. The metabolite differences with false discovery rate (FDR) adjusted p-value ≤ 0.01 and fold changes (FC) ≥ 1.5 or ≤ −1.5 were considered significant. The data was visualized with using principle component analysis (PCA) and partial least squares discriminant analysis (PLS-DA) score plots as well as volcano plot generated with MetaboAnalyst 5.0.

A Venn diagram was created using an online tool: http://bioinformatics.psb.ugent.be/webtools/Venn.

The pathway enrichment analysis was conducted using MetaboAnalyst 5.0; The Small Molecule Pathway Database (SMPDB) containing 99 metabolite sets, based on normal human metabolic pathways, was used as referenced library.

## 3. Results

### 3.1. Metabolic dysregulations in triple negative breast cancer cell lines

We investigated metabolic differences between 5 triple negative breast cancer cell lines (BT549, HCC1143, MDA-MB-231, MDA-MB-436, and MDA-MB-468) and normal breast cell line hTERT-HME1 using untargeted broad metabolic profiling. A total of 479 metabolites were quantified across eight primary pathways related to metabolism of amino acids, carbohydrates, cofactors and vitamins, energy, lipids, nucleotides, peptides, and xenobiotics in the samples. The clear separation, which can be seen between TNBC and control cells, on the principal component analysis (PCA) score plots (**Supplementary Figure 1, Figure 1A**), suggest strong metabolic differences between normal and cancer cells.

**Figure 1.**
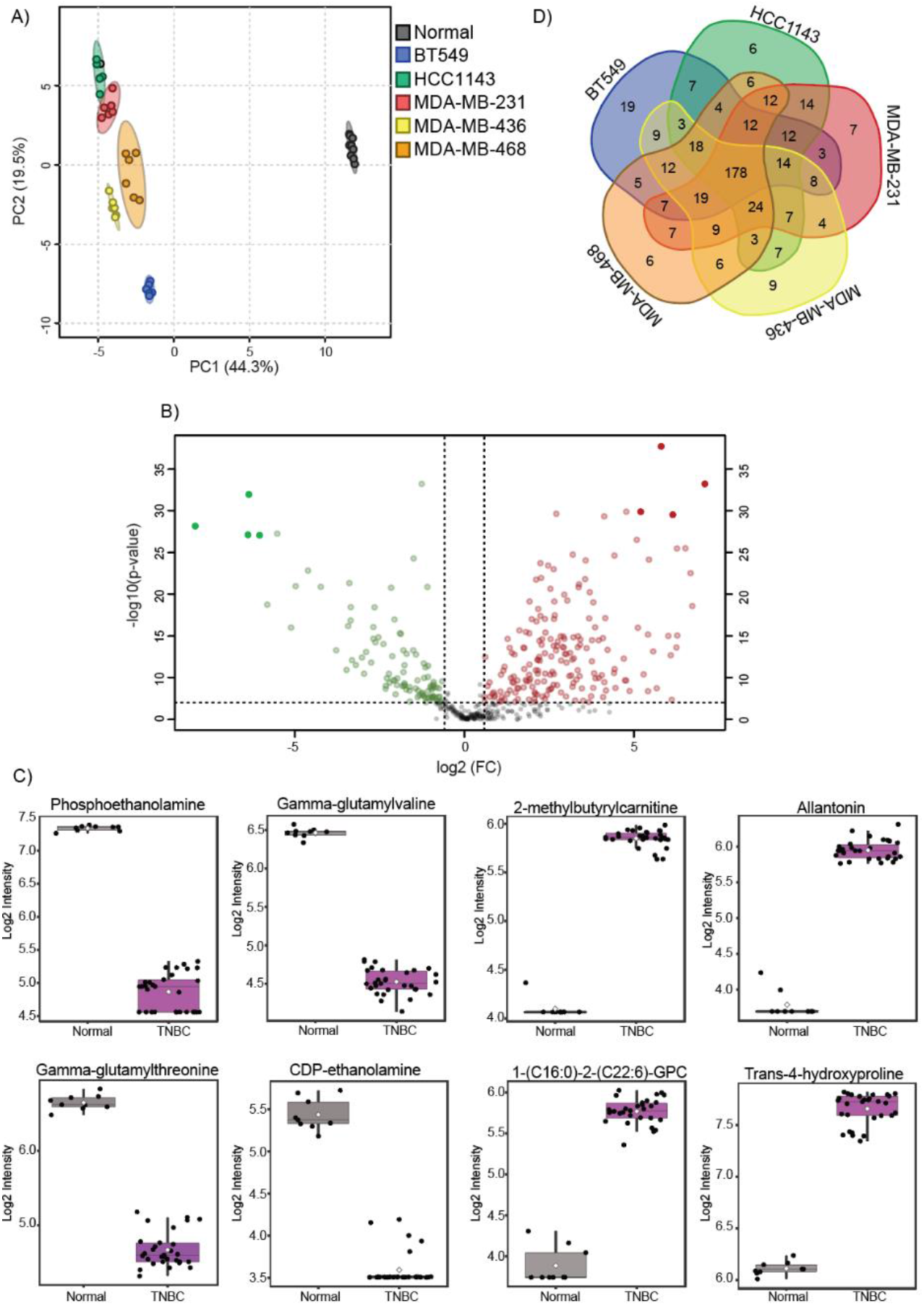
Metabolic signatures of TNBC cell lines. **A)** PCA analysis reveals metabolic differences between normal and TNBC cell lines. **B)** Volcano plot of metabolic features that significantly (with FDR p-value < 0.01 and fold-change ≥1.5 or ≤ −1.5) differ between normal and TNBC cell lines. **C)** Box plots of metabolic features that show the strongest differences between normal and TNBC cell lines. **D)** Venn diagram of similarities and differences across TNBC cell lines relative to normal cells.

The separation between different cancer cell lines is also confirming cancer cell line specific metabolic fingerprints. Out of 479 detected metabolites 291 showed significant, FDR adjusted (FDR < 0.01) and simultaneously fold-change (FC) > 1.5 or < −1.5 differences between normal and TNBC cells across various pathways (**Supplementary Table 1**). The molecules predominantly involved in the metabolism of lipids (117 molecules), amino acids (94 molecules), nucleotides (21 molecules), carbohydrates (18 molecules), and cofactors and vitamins (16 molecules) differentiate the TNBC cell lines from normal cells. Among lipids that were significantly differently regulated we found mainly glycerophospholipids (lysophosphatidylcholines, phosphatidylcholines, and phosphatidylethanolamines), sphingolipids and fatty acids; the amino acids (identified as significantly altered), including branched chain amino acids (BCAA), aromatic amino acids (AAA), methionine and glutathione metabolites. The volcano plot (**Figure 1B**) highlights metabolites with FDR p-value ≤ 0.01 and fold-change (FC) ≥1.5 or ≤ −1.5. The top 4 hits showing up or down regulation in TNBC cells were selected for visualization (**Figure 1C**).

Next, to identify similarities across cancer cell lines as well as their unique metabolic fingerprints we investigated the differences between each cancer cell line and the normal cell line. This comparison revealed 178 metabolites showing common alterations across all examined cancer cell lines as well as unique, cancer cell line specific metabolic features (**Figure 1D**). The largest number of unique metabolites differentiating normal and breast cancer cell lines was identified for BT549 cell line, which is in accordance with PCA showing greatest separation of this cell line on the score plot. This data provides an overview on metabolic features differentiating TNBC cell lines from normal cells as well as emphasizes cancer cell line metabolic individuality.

### 3.2. Triple negative breast cancer cell lines with low and high metastatic potential exhibit different metabolic profiles

A previous study defining metastatic potential of 500 different human cancer cell lines *in vivo* characterized BT549 and HCC1143 as cell lines with low metastatic potential, whereas MDA-MB-231, MDA-MB-436 and MDA-MB-468 were shown to constitute cell lines with high metastatic potential [32]. Thus, we followed this categorization and investigated whether those cell lines possess diverse metabolic program *in vitro*.

First, we monitored the abundance of protein markers of EMT (E-cadherin, N-cadherin, and vimentin) [21] and other proteins which were previously characterized as key components of metastatic cascade (MMP2, TWIST2 and p53) [38–40]. We detected E-cadherin, an important in maintaining epithelial phenotype, in 2 cell lines including HCC1143 and MDA-MB-468 (**Figure 2A**).

**Figure 2.**
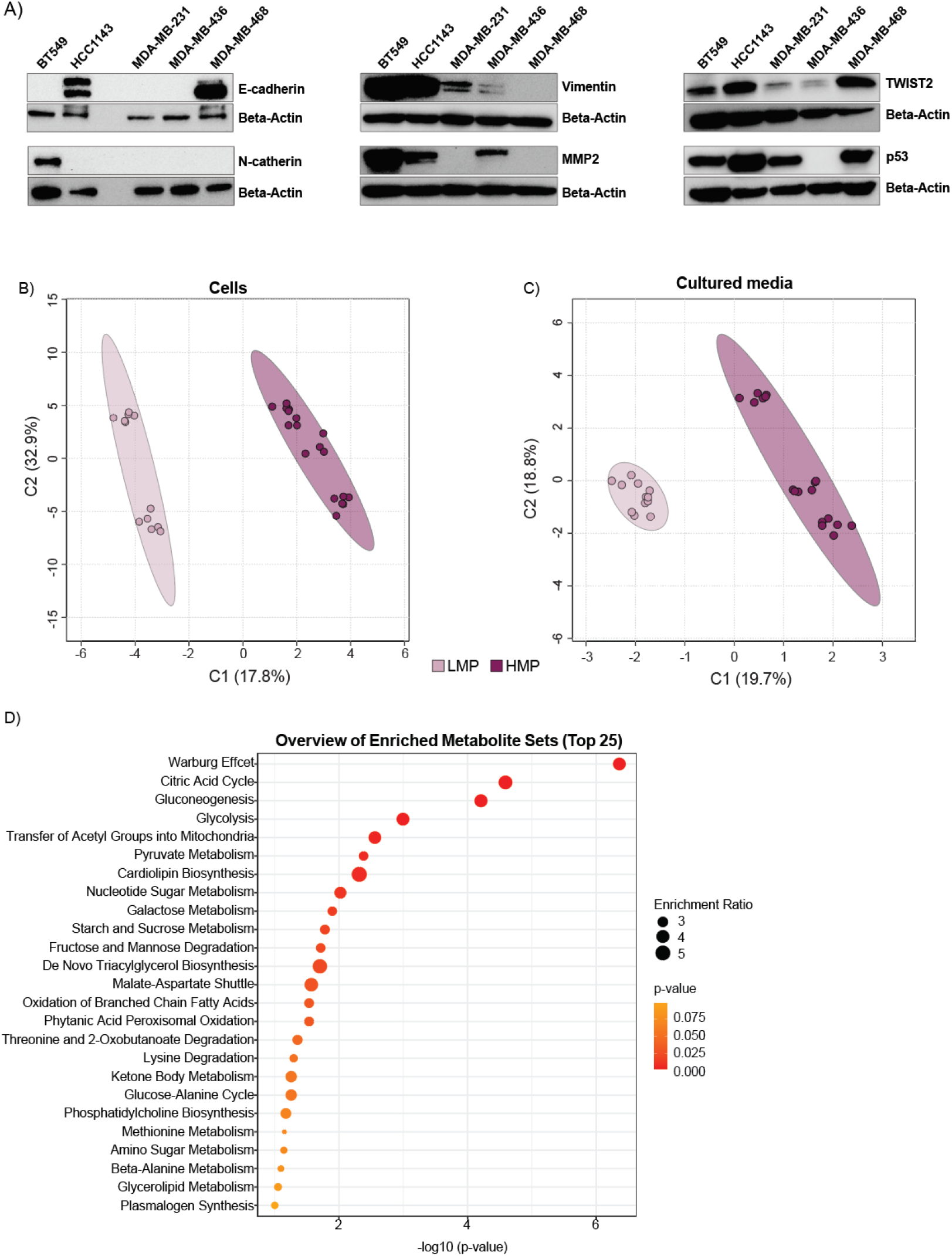
The LMP and HMP cell lines differ metabolically, and this is independent of the level of canonical markers of EMT. **A)** Protein levels of EMT markers and other molecules involved in metastatic cascade. **B)** and **C)** partial least squares discriminant analysis (PLS-DA) score plot conducted on metabolites detected in cells and growth media. **D)** Pathway enrichment analysis plot of cellular metabolism.

N-cadherin, was detected only in BT549 cell line and vimentin, in four (BT549, HCC1143, MDA-MB-231, and MDA-MB-436) out of five examined cell lines (**Figure 2A**). MMP2, was detected in BT549, HCC1143 and MDA-MB-436 and TWIST2 showed the highest expression in MDA-MB-468 (**Figure 2A**). The cell line MDA-MB-436 was lacking expression of p53. This data show that selected cell lines strongly differ in canonical EMT markers as well as components involved in metastatic cascade, and those differences were not reflecting on their metastatic potential.

Next, we investigated whether the selected cell lines displaying different metastatic potential *in vivo* exhibit already distinct metabolic phenotypes *in vitro.* To that end, we conducted metabolic profiling of both cells as well as growth media. The partial least squares discriminant analysis (PLS-DA) on metabolite profiles from both the cells as well as growth media of all TNBC cell lines revealed separation between LMP and HMP in cells (Figure 2B) and growth media (Figure 2C), which suggest metabolic differences between LMP and HMP. We further tested for the metabolites showing FDR (p-value ≤ 0.01) significant differences and the FC ≥ 1.5 or ≤ −1.5 between LMP and HMP. We 92 metabolites in cells (**Table 1**) and 22 metabolites in growth media (**Table 2**), distinguishing LMP from HMP.

**Table 1.**
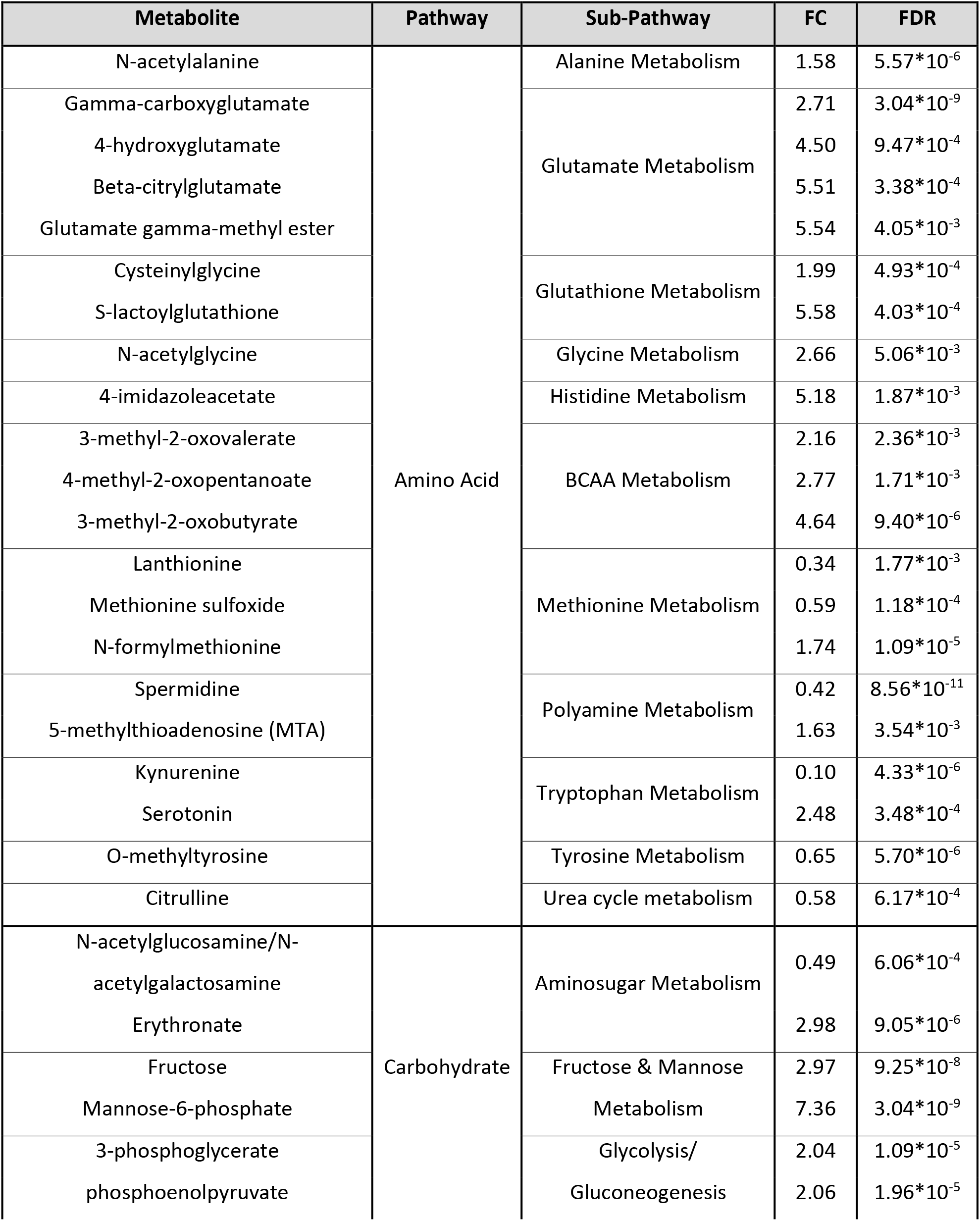
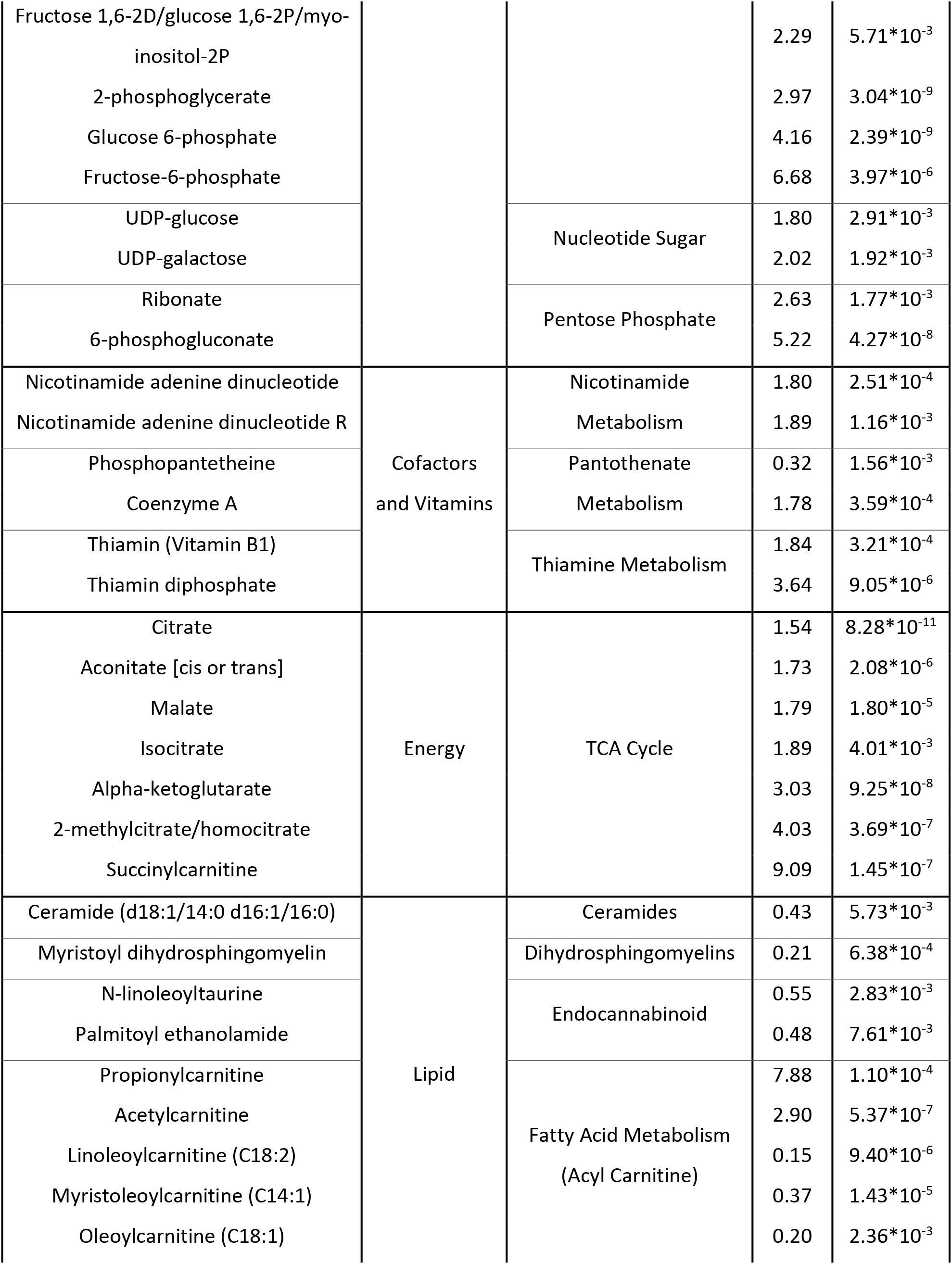
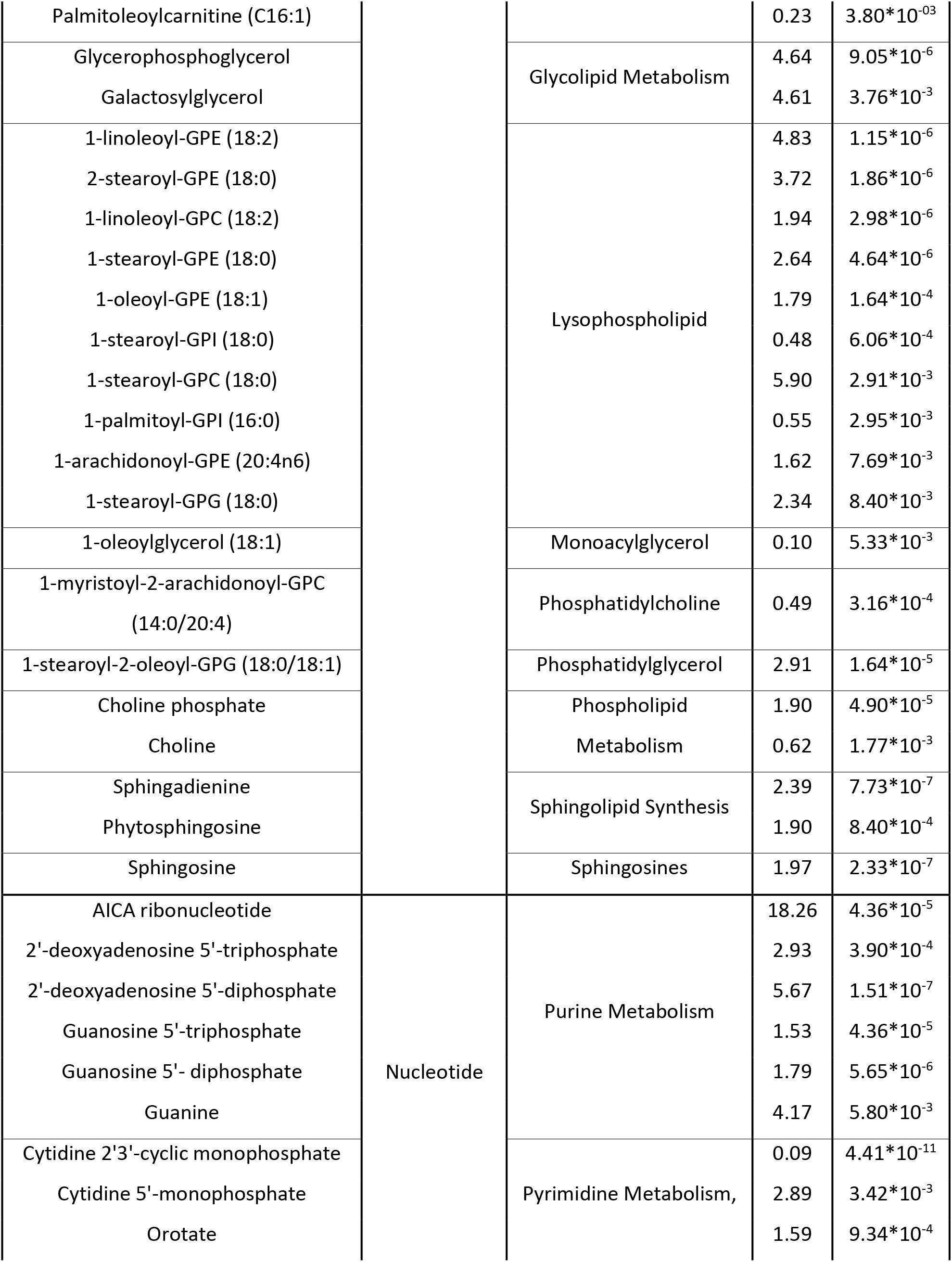
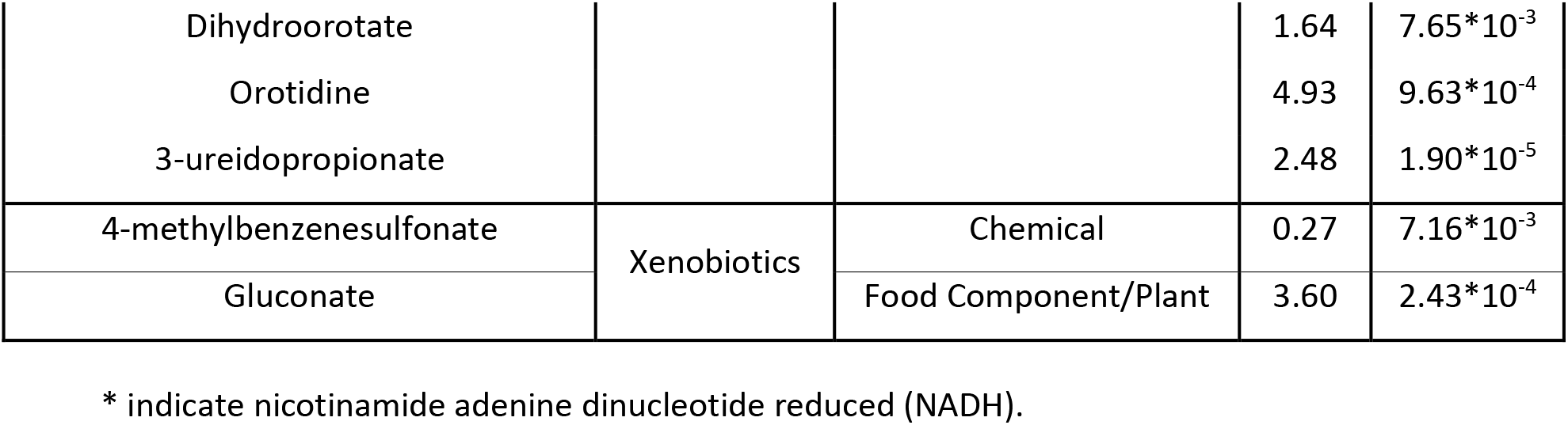
List of cellular metabolites significantly differentiating cell lines harboring HMP from LMP.

**Table 2.** List of growth medium metabolites significantly differentiating cell lines harboring HMP from LMP.

| Metabolite | Pathway | Sub-Pathway | FC | FDR |
| --- | --- | --- | --- | --- |
| Guanidinoacetate | Amino Acid | Creatine Metabolism | 0.39 | 1.35*10<sup>-3</sup> |
| 2-hydroxy-3-methylvalerate |  | BCAA Metabolism | 3.62 | 1.19*10<sup>-13</sup> |
| Alpha-hydroxyisocaproate |  |  | 4.08 | 2.30*10<sup>-6</sup> |
| 1-carboxyethylleucine |  |  | 0.66 | 9.39*10<sup>-5</sup> |
| 4-methyl-2-oxopentanoate |  |  | 2.77 | 4.91*10<sup>-4</sup> |
| 3-methyl-2-oxobutyrate |  |  | 2.19 | 5.88*10<sup>-4</sup> |
| 3-methyl-2-oxovalerate |  |  | 2.44 | 6.46*10<sup>-4</sup> |
| Ethylmalonate |  |  | 0.49 | 2.89*10<sup>-3</sup> |
| Cystathionine |  | Methionine Metabolism | 0.57 | 1.92*10<sup>-3</sup> |
| Kynurenine |  | Tryptophan Metabolism | 0.12 | 1.51*10<sup>-5</sup> |
| 3-hydroxyhexanoate | Lipid | Fatty Acid, Monohydroxy | 2.67 | 7.38*10<sup>-3</sup> |
| 5-dodecenoate (12:1n7) |  | Medium Chain Fatty Acid | 0.31 | 2.61*10<sup>-5</sup> |
| Choline |  | Phospholipid Metabolism | 0.48 | 7.05*10<sup>-4</sup> |
| Docosahexaenoate (22:6n3) |  | Polyunsaturated Fatty Acid (n3 and n6) | 0.34 | 2.10*10<sup>-7</sup> |
| Eicosapentaenoate (20:5n3) |  |  | 0.52 | 4.35*10<sup>-3</sup> |
| Thymine | Nucleotide | Pyrimidine Metabolism | 0.33 | 9.39*10<sup>-5</sup> |
| Uracil |  |  | 0.49 | 1.10*10<sup>-5</sup> |
| 3-ureidopropionate |  |  | 2.39 | 1.09*10<sup>-4</sup> |
| Phenylacetyl glycine | Peptide | Acetylated Peptides | 0.54 | 9.80*10<sup>-3</sup> |
| Gamma-glutamyl glycine |  | Gamma-glutamyl Amino Acid | 1.67 | 5.53*10<sup>-3</sup> |
| p-aminobenzoate | Xenobiotics | Benzoate Metabolism | 0.45 | 3.50*10<sup>-3</sup> |
| 3-hydroxyhippurate |  |  | 1.72 | 6.75*10<sup>-3</sup> |

Among the 92 cellular metabolites that are significantly different between HMP and LMP cell lines, we found 30 lipids (fatty acids and lysophospholipids), 21 amino acids involved in glutamate, BCAA and methionine metabolism, 14 carbohydrates contributing mainly to glycolysis, 12 nucleotides, 7 TCA cycle metabolites, 6 cofactors and vitamins, and 2 xenobiotics. We conducted enrichment analysis on cellular metabolites using MetaboAnalyst 5.0, and found FDR significant enrichment (p-value < 0.05) in Warburg effect, TCA cycle, gluconeogenesis and glycolysis (**Figure 2D**).

Among 22 metabolites measured in growth media that show significant differences between HMP and LMP cell lines 10 were amino acids including 7 molecules involved in BCAA metabolism, 5 lipids, 3 nucleotides and 2 xenobiotics. The metabolic signatures of HMP cell lines identified in media were not showing significant any enrichment.

Taken together, these findings indicate that cell lines with LMP and HMP differ metabolically *in vitro,* and those differences are independent of the EMT markers.

### 3.2. Metabolic pathways contributing to metastatic potential of cancer cells

We constructed the metabolic pathway based on the molecules showing significant differences between HMP and LMP cell lines (**Supplementary Figure 2**). The metabolic signatures differentiating HMP from LMP cell lines focuses around three main pathways namely glycolysis, TCA cycle and lipid metabolism. In addition, we identified a significant increase in the levels of 2-hydroxy-3-methylvalerate (p-value =1.19*10^-13^; FC = 3.62), alpha-hydroxyisocaproate (p-value =2.30*10^-6^; FC = 4.08) in growth media and 4-methyl-2-oxopentanoate (media: p-value =4.91*10^-4^; FC = 2.77; cell: p-value =1.71*10^-3^; FC = 2.77), 3-methyl-2-oxobutyrate (media: p-value =5.88*10^-4^; FC = 2.19; cell: p-value =9.40*10^-6^; FC = 4.64), and 3-methyl-2-oxovalerate (media: p-value =6.46*10^-4^; FC = 2.44; cell: p-value =2.36*10^-3^; FC = 2.16) in both growth media and cells (**Supplementary Figure 3**). All those mentioned metabolites are products of BCAA catabolic pathway; the largest differences observed in the levels of 2-hydroxy-3-methylvalerate, alpha-hydroxyisocaproate in media and 3-methyl-2-oxobutyrate indicate accelerated catabolism of leucine and valine, respectively. Elevated levels of those metabolites in growth media suggest their release by the HMP cells.

In HMP cells, we also observed the elevated levels of cellular gamma-carboxyglutamate (p-value =3.04*10^-9^; FC = 2.71) and 4-hydroxyglutamate (p-value =9.47*10^-4^; FC = 4.50), which are products of glutamate metabolism; the glutamate and glutamine levels were not different between cell lines with distinct metastatic potential (**Supplementary Figure 4 A**). Similarly, differences in the levels of products of tryptophane metabolism, namely kynurenine in both media (p-value =9.47*10^-4^; FC = 4.50) and cells (p-value =9.47*10^-4^; FC = 4.50) as well as cellular serotonin level (p-value =9.47*10^-4^; FC = 4.50) but not tryptophan, were found significantly different between HMP and LMP (**Supplementary Figure 4 B**). Additionally, lower cellular level of spermidine (p-value =8.56*10^-11^; FC = 0.42) was found in HMP in comparison to LMP without changes in other polyamines (**Supplementary Figure 4 C**).

#### 3.2.2. Elevated glycolysis is a signature of cell lines harboring high metastatic potential

Elevated glucose metabolism is a well-known hallmark of cancer [41], which was shown to be even more dysregulated in metastatic cells [42]. We found enrichment in glucose metabolism across metabolites differentiating cell lines with HMP and LMP (**Figure 2D**). We further focused on the molecules involved in carbohydrate metabolism showing significant differences between HMP and LMP cell lines (**Figure 3**).

**Figure 3.**
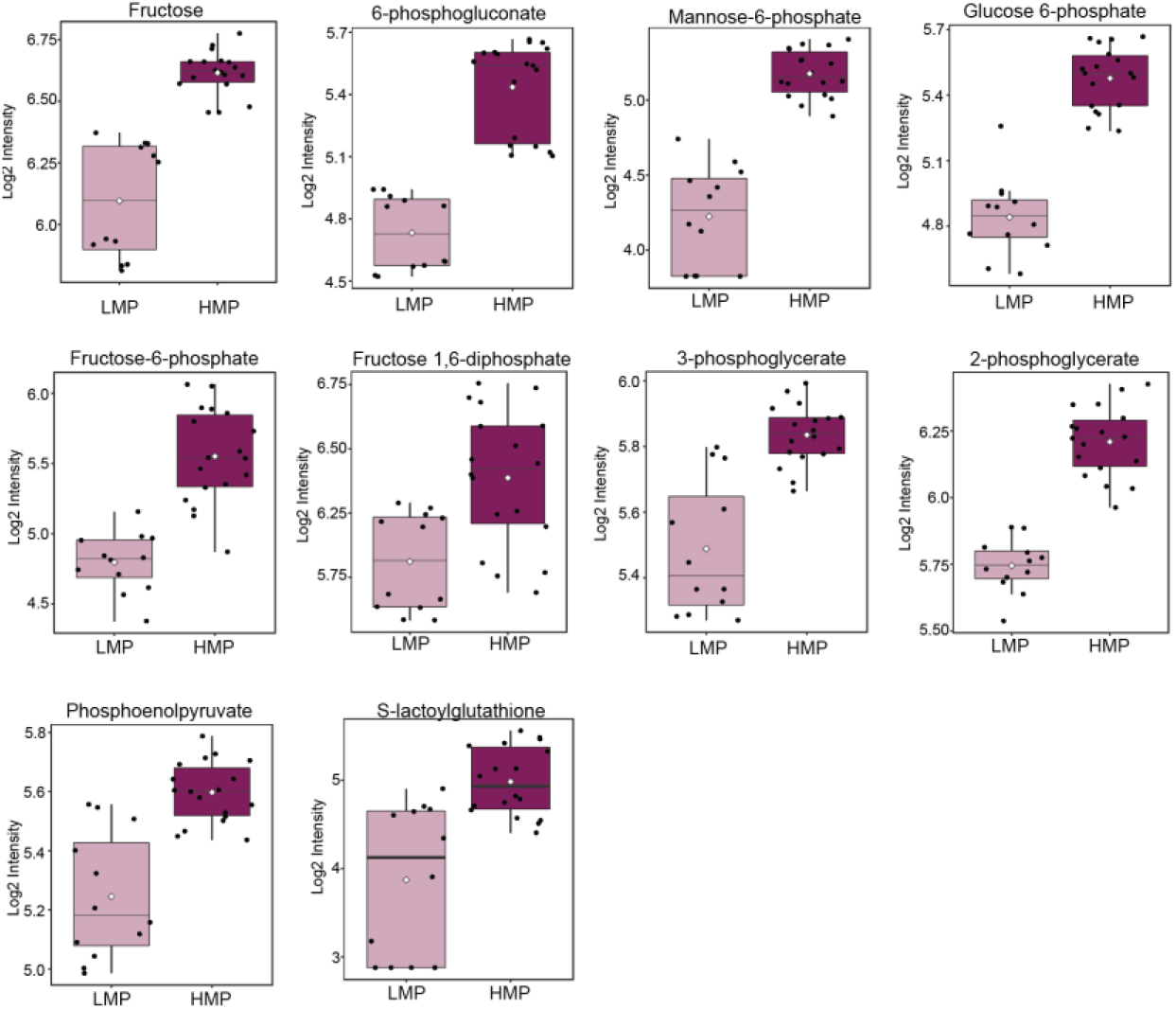
The cell lines with HMP manifest increased glycolysis. The data is presented in the form of box plots. Light purple indicates cells with LMP; dark purple indicates cells with HMP.

The molecules involved in glycolysis including glucose 6-phosphate, fructose-6-phosphate, fructose 1,6-diphosphate, 3-phosphoglycerate, 2-phosphoglycerate, and phosphoenolpyruvate (PEP) (**Figure 3,** and **Table 1**) were all elevated in cell lines harboring HMP. The levels of lactate and pyruvate were not significantly different between cell lines with distinct metastatic potential, but the level of S-lactoylglutathione, which is a part of pyruvate pathway and can be metabolized into lactate, was significantly elevated in HMP cell lines. The glycolytic pathway is interconnected with pentose phosphate metabolism as well as contributes to purine and pyrimidine synthesis [43]. We observed elevated levels in HMP of two molecules (6-phosphogluconate and ribonate) of pentose phosphate pathway (**Figure 3**, and **Table 1**) as well as elevated levels of purines containing adenine (2’-deoxyadenosine 5’-diphosphate (p-value =1.51*10^-7^; FC = 5.67) and 2’-deoxyadenosine 5’-triphosphate (p-value =3.90*10^-4^; FC = 2.93)) and guanine (guanine (p-value =5.80*10^-3^; FC = 4.17), guanosine 5’-diphosphate (p-value =5.65*10^-6^; FC = 1.79) and guanosine 5’-triphosphate (p-value =4.36*10^-5^; FC = 1.53)) and AICA ribonucleotide (p-value =4.36*10^-5^; FC = 18.26) as well as pyrimidines containing orotate (dihydroorotate (p-value =7.65*10^-3^; FC = 1.64) and orotate (p-value =9.34*10^-4^; FC = 1.59)) and cytidine (cytidine 5’-monophosphate (p-value =3.42*10^-3^; FC = 2.89)). The level of cytidine 2’3’-cyclic monophosphate was significantly (p-value =4.41*10^-11^; FC = 0.09) decreased in cell lines with HMP in comparison with those of LMP. The HMP cell lines displayed elevated levels of fructose and manose-6-phospahete in comparison with LMP cell lines (**Figure 3**, and **Table 1**). Thus, it could be reasoned that cell lines harboring HMP exhibit elevated glycolysis, potentially to supply nucleotide synthesis.

#### 3.2.3. Upregulated citrate metabolism but not entire TCA cycle is a hallmark of cell lines with HMP

Dysregulated metabolism of TCA cycle molecules, including succinate, fumarate, alpha-ketoglutarate, 2-hydroxyglutarate and citrate, was previously attributed to metastatic cells [23]. Our enrichment analysis suggested that TCA cycle metabolism is enhanced in TNBC cell lines with HMP (**Figure 2 D**). We observed that mainly citrate and the components of citrate metabolism, including aconitate [cis or trans], beta-citrylglutamate, isocitrate and alpha-ketoglutarate, were significantly elevated in cell lines harboring HMP (**Figure 4** and **Table 1**). The levels of succinyl-CoA, succinate and fumarate were not significantly different between HMP and LMP cell lines. The malate level was higher in HMP in comparison to LMP cell lines. We have also found increased levels of succinyl carnitine and propionyl carnitine (**Figure 4** and **Table 1**) which can contribute to TCA cycle on the succinyl-CoA level. Interestingly, levels of thiamine and thiamine diphosphate, which are critical for the activity of TCA cycle enzymes including pyruvate dehydrogenase (PDH) and alpha-ketoglutarate dehydrogenase (α- KGDH) [44], were significantly elevated in HMP cell lines (**Figure 4** and **Table 1**). Taken together, the citrate metabolism, rather than entire TCA cycle, is upregulated in cell lines with HMP.

**Figure 4.**
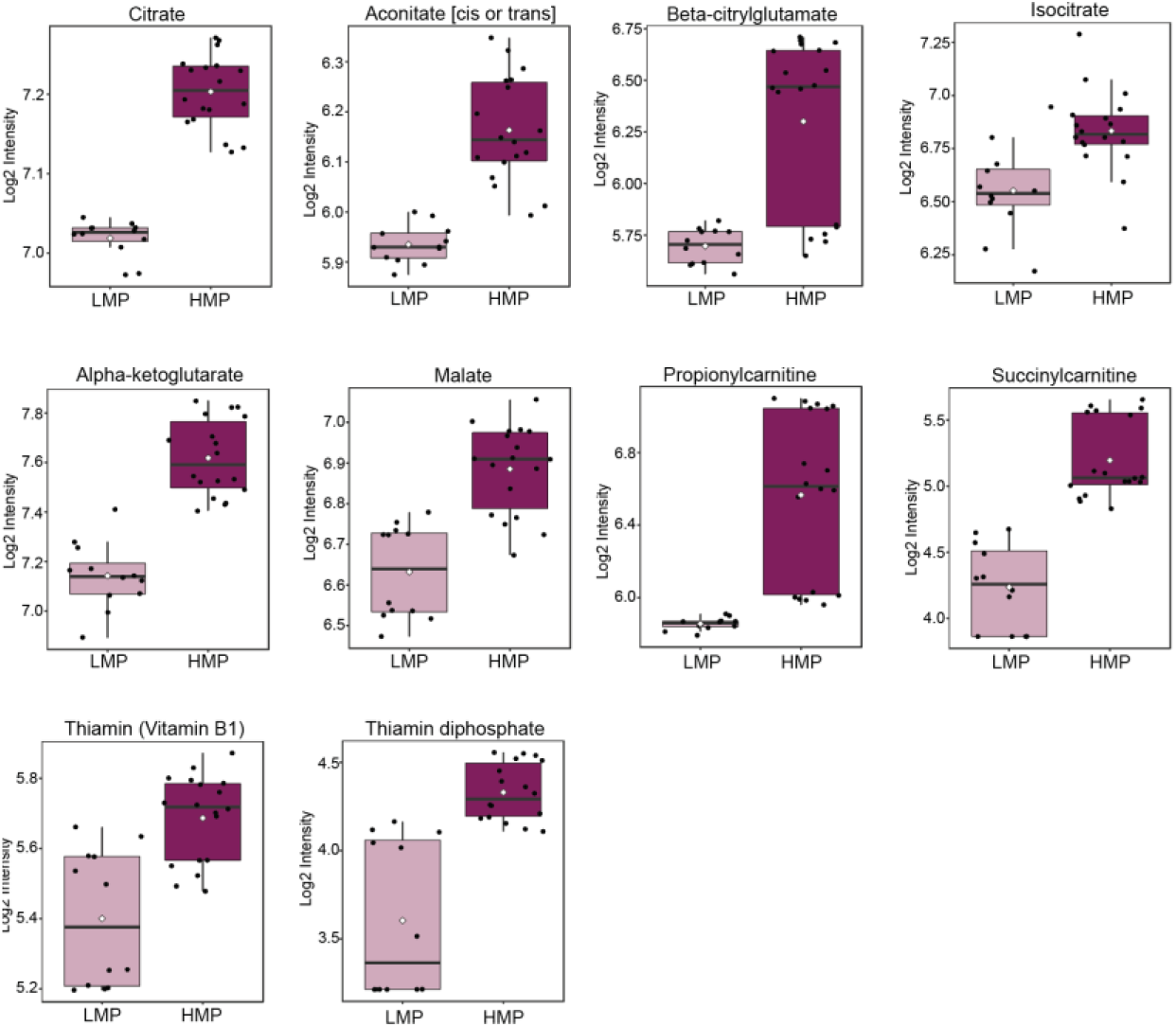
The cell lines harboring HMP exhibit enhanced citrate metabolism. The data is presented in the form of box plots. Light purple indicates cells with LMP; dark purple indicates cells with HMP.

#### 3.2.4. Dysregulated lipid metabolism is a feature of cell lines with HMP

We identified 30 different molecules involved in lipid metabolism, which significantly differentiate HMP from LMP cell lines (**Table 1**). To provide further insight regarding their potential contribution to cancer cell metastatic we have analyzed them in the context of a pathway (**Supplementary Figure 2**). The levels of lysophospholipids were differently regulated between LMP and HMP namely lysophosphatidylinositols were significantly decreased whereas lysophosphatidylcholines and lysophosphatidylethanolamines were significantly elevated in cell lines with HMP in comparison to the one with LMP (**Figure 5** and **Supplementary Figure 2**). The levels of choline were decreased whereas phosphocholine levels were elevated in HMP cell lines (**Figure 5**). Out of 43 measured glycerophospholipids only two (including elevation in HMP 1-stearoyl-2-oleoyl-GPG (18:0/18:1) and decrease in HMP 1-myristoyl-2-arachidonoyl-GPC (14:0/20:4)) were showing significant differences between HMP and LMP cell lines (**Figure 5**). We did not observe any differences in the free fatty acid levels between LMP and HMP but found significantly lower levels of four acylcarnitines (linoleoylcarnitine (C18:2), oleoylcarnitine (C18:1), palmitoleoylcarnitine (C16:1), and myristoleoylcarnitine (C14:1)), in cell lines harboring HMP (**Figure 5** and **Table 1**). The acylcarnitines are required to transport free fatty acids across mitochondrial membrane for beta-oxidation. The lower level of acylcarnitines could suggest decrease in beta-oxidation in HMP cell lines and incorporation of fatty acids into lysophospholipids which were significantly elevated (**Figure 5** and **Table 1**). The level of acetylcarnitine was significantly elevated in HMP cell lines. We also identified alteration in sphingolipid metabolism; the levels of sphingosine and phytosphingosine were elevated whereas sphingomyelins decreased in the cell lines with HMP. Taken together, this data indicates that cancer cell lines harboring distinct metastatic potential activate different programs of lipid metabolism. The lipid dysregulation in HMP manifests in increased levels of glycerophospholipids and acetylcarnitine and decreased levels of acylcarnitines further suggesting potential enhancement in lipid synthesis.

**Figure 4.**
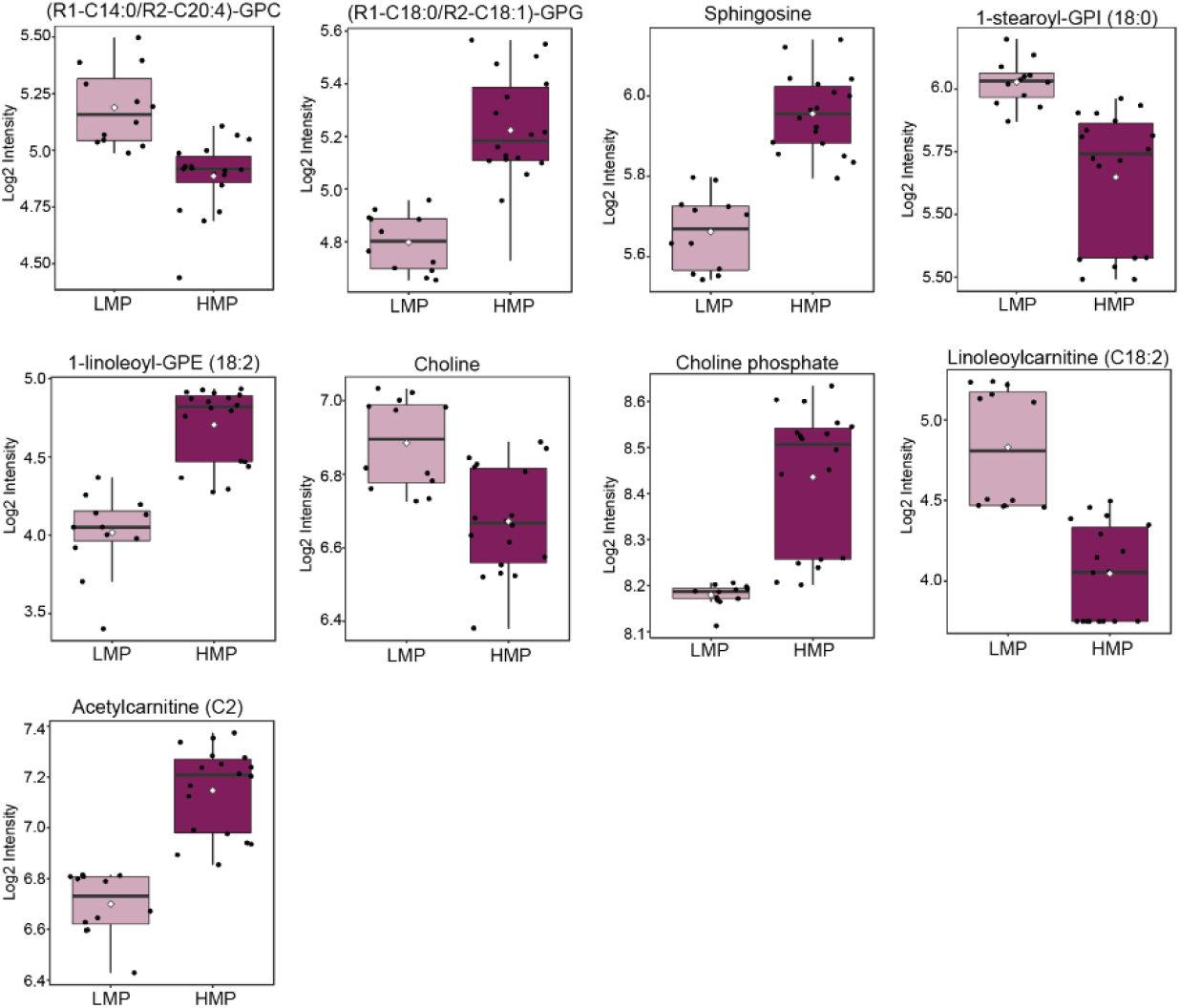
The components of lipid metabolism differentiate HMP from LMP cell lines. Box plots showing examples of alterations in molecules involved in various lipid metabolism pathways. Light purple indicates cells with LMP; Dark purple indicate cells with HMP.

## 4. Discussion

Dysregulated metabolism plays a vital role in cancer cell progression and metastasis [22,23,25,26,45–49]. In this study, we have shown that TNBC cell lines differentiating in their metastatic potential *in vivo* exhibit different metabolic profile already *in vitro,* and those differences are independent of the abundance of canonical EMT markers. The cell line MDA-MB-468 reported as with HMP in vivo, was not expressing any of the canonical markers of EMT. Nevertheless, to undergo EMT this cell line requires exposure to epidermal growth factor (EGF) [41], thus in vitro abundance of EMT markers was not expected for this particular cell line. Cell lines harboring HMP displayed enrichment in glycolysis and TCA cycle, as well as dysregulated metabolism of lipids. The elevated levels of products of BCAA catabolism in cells as well as growth media suggested accelerated BCAA catabolism in TNBC cell lines with HMP. Additionally, we found significant differences in the levels of gamma-carboxyglutamate, 4-hydroxyglutamate, kynurenine, serotonin, and spermidine between HMP and LMP cell lines.

Elevated glycolysis was previously described as a feature of metastatic cells, which supports cancer cell survival under energy deficient conditions occurring after cell dissociation from the primary tumor and establishment of metastatic niche [47]. Moreover, enhanced glycolysis was shown to support release of exosome by metastasizing cells, which is a crucial step in the metastatic cascade, and thus linking dysregulated glucose metabolism with cancer cell ability to extravasate at distant premetastatic niche [48]. Therefore, the enhanced glycolytic program in the cells with HMP, observed in our study, is in concordance with previous reports and further pinpoints metabolic advantage of HMP over LMP cell lines. Elevated glycolysis in cancer cells frequently results in accumulation of methylglyoxal, which is further metabolized to S-lactoylglutathione by glyoxalase 1 (GLO1) and subsequently to lactate in the reaction catalyzed by glyoxalase 2 (GLO2) [50]. The elevated expression of GLO1 was found in basal TNBC and was shown to be essential for the survival of breast cancer stem cells [51]. The elevated level of S-lactoylglutathione observed in our study in cell lines with HMP further underscores enhanced glycolytic program in those cells as well as suggests increased expression of GLO1. Importantly, S-lactoylglutathione could serve as reservoir of lactate, shown to be a key player during metastasis by stimulating angiogenesis and increasing extracellular acidification to evade the immune system [52], which could be considered as further metabolic advantages of HMP cell lines.

Elevated glycolysis in cancer cells is frequently linked with enhanced lactate synthesis, however differences in the lactate level between cell lines harboring HMP and LMP was not found. Nevertheless, we found increased level of citrate and molecules involved in citrate metabolism, which could suggest higher glucose contribution to TCA cycle in HMP cell lines. The greater flux of glucose into TCA cycle was recently reported in metastatic colorectal cancer cell lines [49]. The enhanced glycolysis along with intracellular accumulation of citrate were shown to enhance TNBC cell invasion and metastasis via AKT/ERK signaling pathway [46]. In our study in addition to elevated citrate level we found increased levels of other molecules involved in citrate metabolism including aconitate [cis or trans], isocitrate, alpha-ketoglutarate as well as beta-citrylglutamate further highlighting potential importance of this pathway in governing the metastatic cascade. Importantly, beta-citrylglutamate was shown as an activator of aconitase, which catalyzes isocitrate formation [53]. These results indicate that metabolism of citrate rather than dysregulation of the entire TCA cycle predispose metastatic potential in TNBC cell lines. Importantly, the elevated citrate level is an indicator of energy excess and cell readiness for fatty acid synthesis. Citrate is cleaved by ATP citrate lyase and the achieved oxaloacetate reenters into TCA cycle in the form of malate [54]. The elevated citrate metabolism along with increased malate level in HMP cell lines further suggests enhanced fatty acid and other lipid synthesis in those cells. In concordance, we have observed decrease in the acylcarnitine levels, which are indicators of fatty acid catabolism, as well as an increase in glycerophospholipid levels in HMP cell lines. This observation suggests that enhanced citrate metabolism contributes to accelerated lipid synthesis in HMP cell lines, which was previously linked with tumor progression [55].

Additionally, we observed an increase in the catabolic pathway of BCAA in HMP cell lines manifested by the accumulation of their products of catabolism in both cell and growth media. The products of leucine catabolism, namely 2-hydroxy-3-methylvalerateand alpha-hydroxyisocaproate, displayed the greatest accumulation in HMP cell lines in comparison with LMP cell lines. The increased BCAA catabolism could contribute to enhanced energy generation and biomass production as well as promotion of mTOR signaling, which is a known cancer cell molecular pathway [56]. Furthermore, it was shown that inhibition of leucine uptake suppresses mTOR signaling and promotes apoptosis in breast cancer cell lines [57]. Moreover, the enhanced activity of branched-chain α-keto acid dehydrogenase kinase (BCKDK), the key enzyme of BCAAs metabolism, was shown to promote migration, invasion and EMT of colorectal cancer [58]. Therefore, it could be reasoned that increased BCAA catabolism observed in HMP cell lines contribute to their metabolic advantage which empowers their metastatic potential.

Noteworthy, the levels of thiamine and thiamine diphosphate, which are critical for activity of enzymes involved in TCA cycle, pentose phosphate and BCAA metabolism, were elevated in cell lines with HMP further underscoring enhanced metabolic potential in those cell lines. The importance of thiamine in cancer cell metabolism was recently suggested [44]. Thus, it could be reasoned that HMP cell lines activate thiamine metabolism to ensure enhanced activity of TCA cycle, pentose phosphate and BCAA metabolism.

Coagulation proteins along with platelets have been shown to promote pro-survival signaling during metastasis [59]. The HMP cell lines displayed elevated levels of γ-carboxyglutamine, which is involved in coagulation cascade; the γ-carboxyglutamic acid residues play an important role in coagulation by governing the activation and binding of circulating blood-clotting enzymes to cell membrane surface [60]. Moreover, the HMP cell lines exhibited an increased level of 2’-deoxyadenosine 5’-triphosphate and 2’-deoxyadenosine 5’-diphosphate, which are molecules of adenine nucleotide metabolism. The role of adenine nucleotides in extravasation was previously suggested and linked with platelet activation by cancer cells [61]. Furthermore, 4-hydroxyglutamate which was also elevated in HMP cell lines, could potentially play a role in platelet activation as this molecule was identified as metabolic marker of preeclampsia, a health condition associated with coagulation and platelet activation [62]. Thus, the cell lines harboring HMP possess metabolic features potentially supporting coagulation and platelet activation, which are important contributors of the metastatic cascade.

## 5. Conclusions

Our study provides new insights into cancer metastasis from the perspective of dysregulated metabolism. The landscape of metabolic dysregulations characterized in our study could serve as a roadmap for identification of treatment strategies targeting cancer cells with enhanced metastatic potential. We identified metabolic advantages of cell lines with high metastatic potential beyond enhanced glycolysis by pinpointing the role of BCAA catabolism as well as molecules supporting coagulation and platelet activation as important contributors to metastatic cascade. A future prospective would be to probe those identified metabolic dysregulations as therapeutic targets.

## Supplementary Materials

### Supplementary Tables

**Supplementary Table 1.** Metabolomics data.

**Supplementary Table 2.** List of metabolites showing FDR significant differences between TNBC and normal cells.

### Supplementary Figures

**Supplementary Figure 1.** Pairwise score plot for top five principal component (PC). Grey - hTERT-HME1; blue - BT549; green - HCC1143; read - MDA-MB-231; yellow - MDA-MB-436; orange - MDA-MB-468.

**Supplementary Figure 2.** Metabolic pathway depicting metabolic differences between HMP and LMP cell lines. Red – metabolites significantly elevated in HMP in comparison with LMP; Green – metabolites significantly decreased in HMP in comparison with LMP; Grey – measured metabolites not significantly different between HMP and LMP; White – metabolite not detected.

**Supplementary Figure 3.** Box plots showing products of BCAA catabolism differentiating significantly HMP and LMP cell lines in **A)** growth medium and **B)** cells. Light and dark purple indicate cell lines with LMP and HMP respectively.

**Supplementary Figure 4.** Box plots showing metabolites differentiating significantly HMP and LMP cell lines involved in **A)** glutamate metabolism; **B)** tryptophane metabolism and **C)** polyamine metabolism. Light and dark purple indicate cell lines with LMP and HMP respectively.

## Author Contributions

Conceptualization, A.H., A.R. and K.S.; methodology, S.K., S.D., G.T., I.A.; formal analysis, A.H.; investigation, A.H., S.K., S.D., G.T., I.A.; writing—original draft preparation, A.H., A.R., K.S., S.K., S.D; writing—review and editing, A.H., A.R., K.S., I.A., S.K., S.D, and G.T.; funding acquisition, A.H, and K.S. All authors have read and agreed to the published version of the manuscript.

