## Supplementary figures and images for "“Metabolic dysregulations of cancer cells with metastatic potential”"

### Supplementary Figure 1.pdf

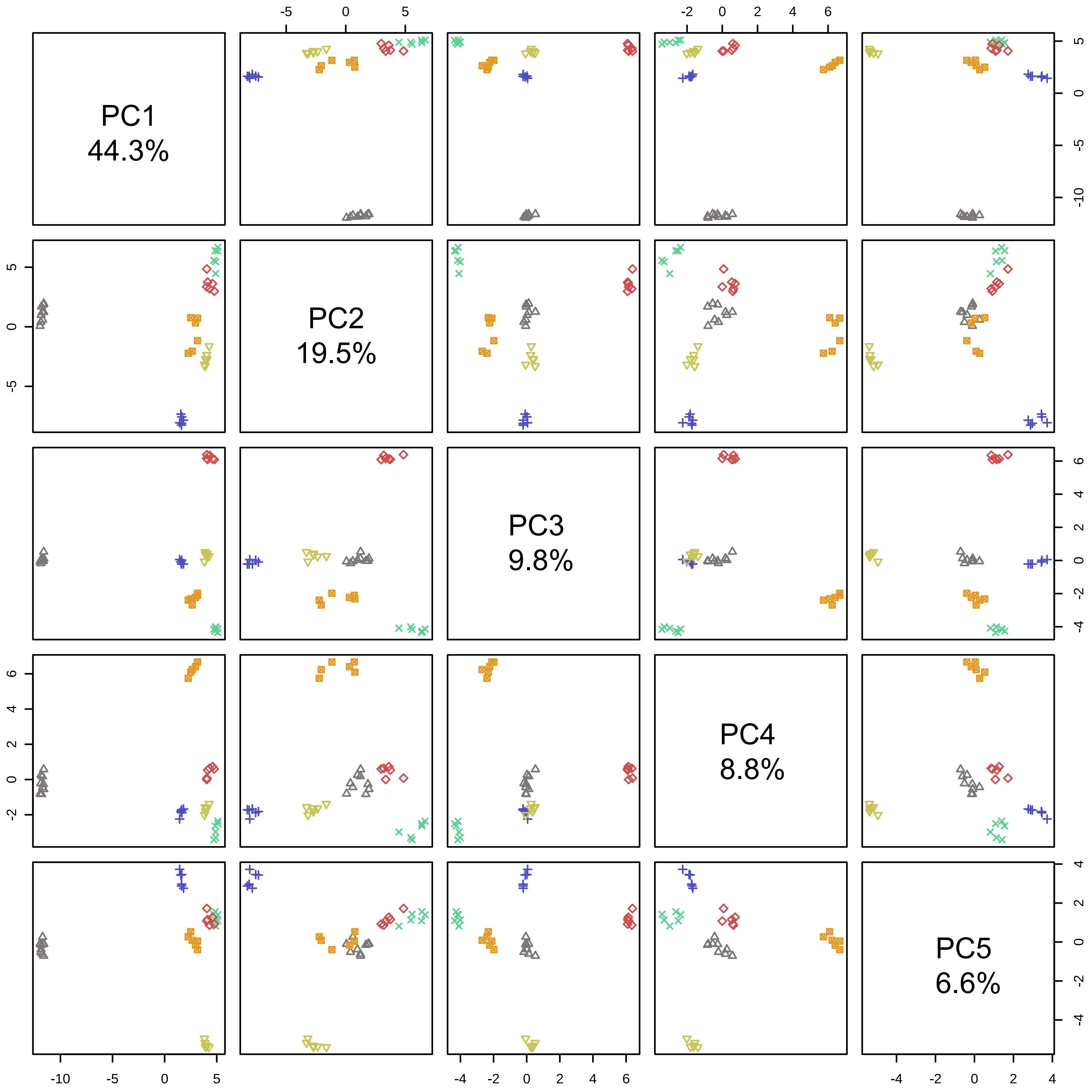

### Supplementary Figure 2.pdf

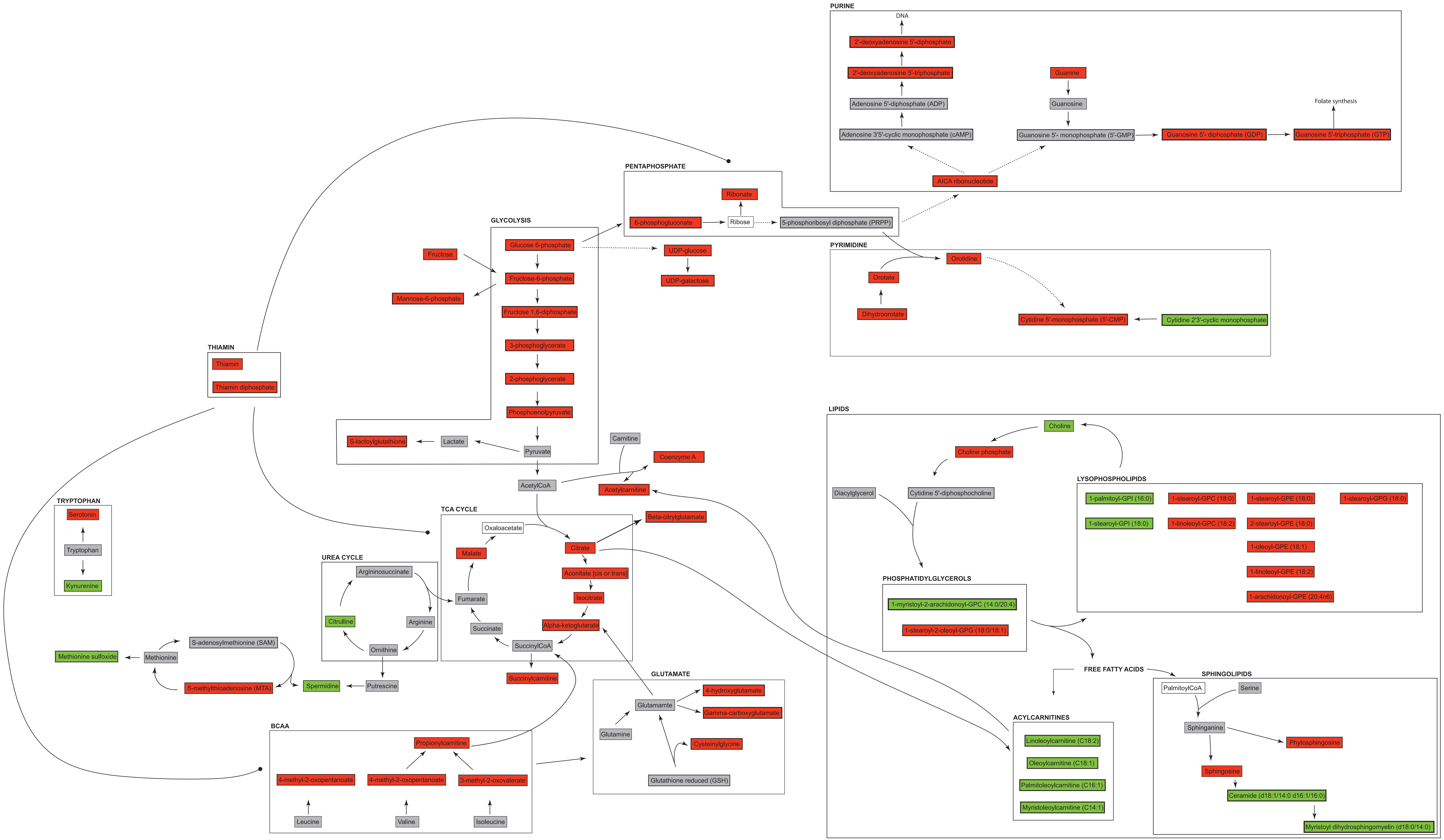

### Supplementary Figure 3.pdf

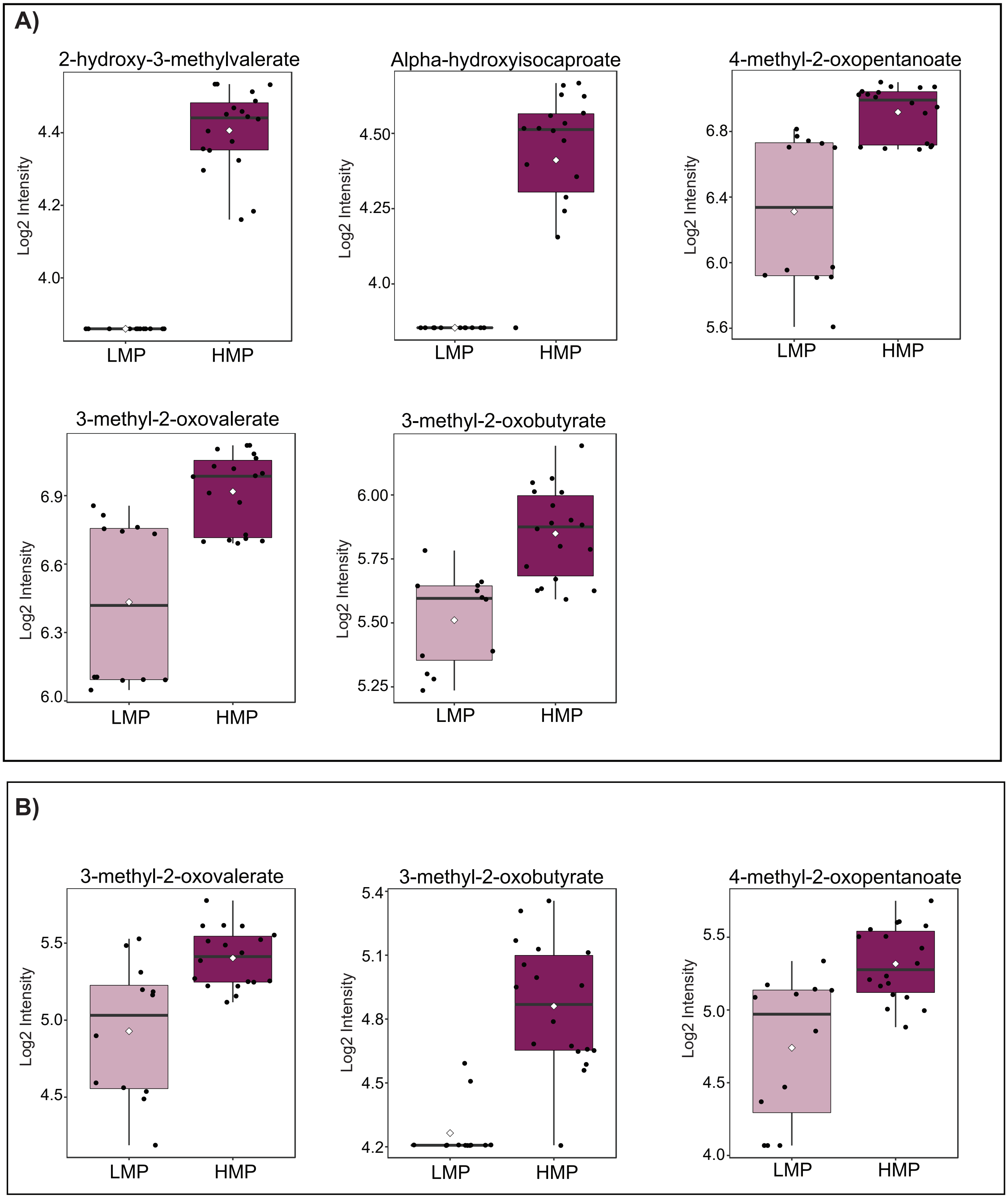

### Supplementary Figure 4.pdf

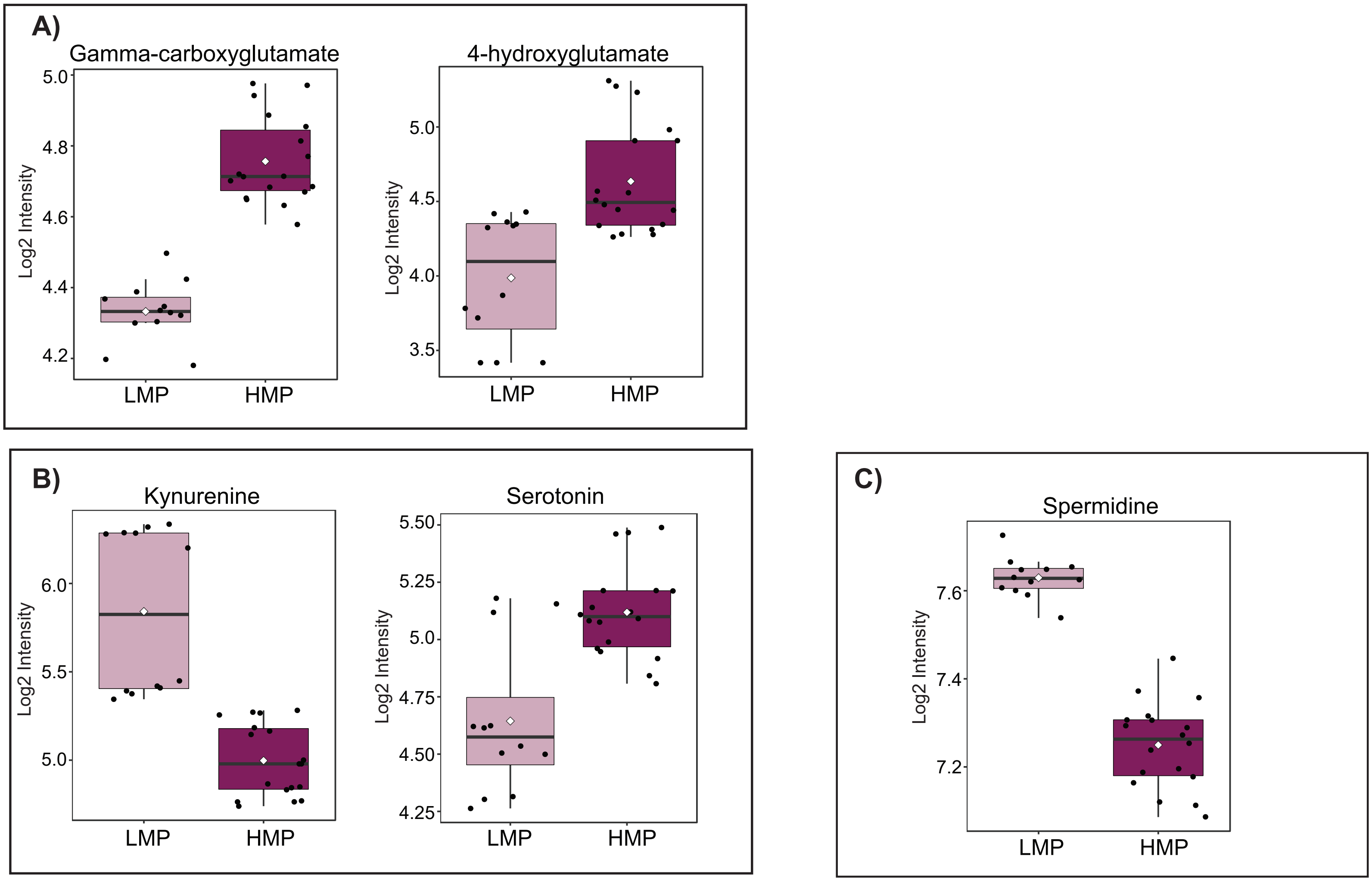
